# Experimental and Computational Investigation of the Structure of Peptide Monolayers on Gold Nanoparticles

**DOI:** 10.1101/083204

**Authors:** Elena Colangelo, Qiubo Chen, Adam M. Davidson, David Paramelle, Michael B. Sullivan, Martin Volk, Raphaël Lévy

## Abstract

The self-assembly and self-organization of small molecules at the surface of nanoparticles constitute a potential route towards the preparation of advanced protein-like nanosystems. However, their structural characterization, critical to the design of bio-nanomaterials with well-defined biophysical and biochemical properties, remains highly challenging. Here, a computational model for peptide-capped gold nanoparticles is developed using experimentally characterized CALNN-and CFGAILSS-capped gold nanoparticles as a benchmark. The structure of CALNN and CFGAILSS monolayers is investigated by both structural biology techniques and molecular dynamics simulations. The calculations reproduce the experimentally observed dependence of the monolayer secondary structure on peptide capping density and on nanoparticle size, thus giving us confidence in the model. Furthermore, the computational results reveal a number of new features of peptide-capped monolayers, including the importance of sulfur movement for the formation of secondary structure motifs, the presence of water close to the gold surface even in tightly packed peptide monolayers, and the existence of extended 2D parallel β-sheet domains in CFGAILSS monolayers. The model developed here provides a predictive tool that may assist in the design of further bio-nanomaterials.

## INTRODUCTION

The electronic and optical properties^1^ of gold nanoparticles (GNPs), together with the possibility of tuning their colloidal and (bio)chemical properties by functionalization with small ligands, such as alkyl thiols and peptides, has led extensive research over the last 20 years, targeted towards potential applications in bioimaging and biomedicine,^2^ sensing and catalysis.^3^ These ligands, through gold-thiol chemistry,^4^ bind to the GNP surface and form compact self-assembled monolayers (SAMs).^5^ Breakthroughs were made in the early ‘90s by studies reporting straightforward synthesis methods^6–8^ to prepare GNPs of desired size, stabilized by alkyl thiols. Ten years later, we introduced peptide self-assembled monolayers on GNPs, starting from the design of CALNN, a pentapeptide developed from general protein folding principles.^9^ We have described peptide-capped GNPs as building-blocks which could potentially be assembled in complex networks and engineered into protein-like objects using a “bottom-up” approach.^10^ Controlling and characterizing the structure of peptide SAMs with molecular details is a fundamental prerequisite to envision building such artificial nanosystems. Attempts at determining the monolayer structure of such systems include the study of Fabris et al., where peptides based on α-aminoisobutyric acid units were found to form 3_10_-helical secondary structure motifs both in solution and at the surface of 1-2 nm GNPs by IR and NMR spectroscopies which also indicated the presence of both intra-and inter-peptide hydrogen bonds.^11^ Rio-Echevarria et al. used circular dichroism to confirm that the helical secondary structure of an undecapeptide was maintained both in solution and at the surface of 1-2 nm GNPs.^12^

While elucidating the structure of peptide monolayers with experimental techniques remains challenging, combining experimental and computational approaches can shed light on the structures with molecular details. Alkyl thiol-capped GNPs have been investigated more extensively than peptide-capped GNPs and different classical force fields (FF) have been reported in the literature describing the interactions of alkyl thiols on gold.^13–16^ The first FF specifically designed for biomolecules on Au(111) surface dates only from 2008.^17^

For such systems, not only the secondary interactions between the amino acids and the gold surface, but also the gold polarization induced by the biomolecule and surrounding water molecules should be taken into account for a compelling description.^18^ The first study using atomistic molecular dynamics (MD) simulations to investigate the structure of alkane thiols on gold nanoparticles as a function of temperature, GNP size and length of the ligand dates from 2007;^19^ investigations of the structure and properties of GNPs capped with a monolayer constituted of two alkane thiols differently organized also followed.^20–22^ However, no MD simulations have been conducted yet to assess the structure of SAMs of peptides on spherical gold nanoparticles while taking into account both the self-organization of peptides and the gold atoms’ polarizability. Examples of simpler descriptions that did not consider such aspects are Duchesne et al.’s work, where MD simulations were used to determine the volume probed by a functional peptide embedded in a mixed SAM peptide monolayer constituting a surface with a curvature appropriate for a 10 nm GNP^23^ and Todorova et al.’s investigation of the effect of TAT concentration, i.e., a cell-penetrating peptide, on the structure and compactness of a CALNN monolayer anchored to 3 nm GNPs built from neutral carbon-type atoms.^24^ Also, Lee et al. looked at the effect that conjugating a single peptide to a GNP had on its structure.^25^

Here, a computational model for peptide-capped gold nanoparticles is presented for which the GolP-CHARMM^26^ force field, parameterized for both Au(100) and Au(111) surfaces and accounting for the dynamic polarization of gold atoms produced by the interactions with the biomolecules and the surrounding environment, was adopted.

In particular, the model is developed using experimentally characterized CALNN-^9^ and CFGAILSS-^27^ capped GNPs as a benchmark and simulations are carried out by means of molecular dynamics. The model’s results for the secondary structure of the monolayers in the simulated systems are compared to those of two independent experimental studies, the first reported here, and the second previously published.^27^ In the first, the effect of capping density on the secondary structure of the CALNN monolayer on GNPs was investigated by Fourier transform infrared (FTIR) spectroscopy. In the second study, the effect of the nanoparticle curvature on the secondary structure and intermolecular interactions of CALNN and CFGAILSS peptide monolayers on GNPs of different sizes was determined by a combination of FTIR and solid-state NMR spectroscopies.^27^ Following validation of the computational model against these experimental studies, detailed insights into the peptides’ structural and dynamic properties are obtained, including the importance of sulfur movement for the formation of secondary structures, the presence of water close to the gold surface even in tightly packed peptide monolayers, and the existence of extended 2D parallel β-sheet domains in CFGAILSS monolayers.

## COMPUTATIONAL MODEL

Molecular dynamics simulations were performed using GROMACS package,^28–31^ version 4.6.5. A combination of the GolP-CHARMM^26^ force field and its predecessor GolP^17^ was adopted to describe the peptides, the gold atoms and the peptide-surface interactions. For the latter, the GolP-CHARMM Lennard Jones parameters for the Au(100) surface were used.

The initial structures for the simulations were prepared as follows: the nanoparticle was modelled as a sphere of the desired size, where the constituting gold atoms, within the GolP-CHARMM force field description, are represented by a real gold atom and a virtual site, called AU and AUC, respectively, where AU is fixed whereas AUC can move (for more details see Supporting Information, section S1.1). AU and AUC are constrained at a distance of 0.07 nm and the mass of each is equal to half the mass of a single gold atom, i.e., 196.97/2 amu. AU and AUC are partially charged, i.e., −0.3 and +0.3, respectively, and thus account for the dynamic polarization of the gold surface. Moreover, the interactions between the peptide atoms and AUC are described only in terms of Coulombic electrostatic interactions, whereas those with AU include both Coulombic and van der Waals interactions; the latter are described by 12-6 Lennard-Jones (LJ) potentials.

The starting coordinates of CALNN (Cys-Ala-Leu-Asn-Asn) and CFGAILSS (Cys-Phe-Gly-Ala-Ile-Leu-Ser-Ser) peptides were created using the Materials Studio program. A linear geometry for the peptide backbone was chosen. The peptides, in zwitterionic form, were oriented with the N-terminus toward the gold nanoparticle surface and the C-terminus located radially away from the nanoparticle. The Cys sulfur atom is deprotonated and bears an extra electron to simulate the strong binding between sulfur atoms and the polarizable gold surface. Overall, the high interaction energy between the charged sulfur atoms and the gold nanoparticle surface is constituted of attractive and repulsive Coulomb terms (S− with AUC and AU, respectively) and a 12-6 LJ potential term (S− with AU, ε of 6.90 kJ/mol and σ of 0.30 nm for AU-S sulfide).^26^ Thus, the cysteine sulfur atoms are responsible for anchoring the peptide on the gold nanoparticle surface, but because they are not directly bound, the computational model developed here has the advantage of accounting for the motion of the sulfur atoms over the nanoparticle surface.^32^ The initial distance between the Cys sulfur atoms and the gold surface was set to 0.25 nm, within the range of 0.22 – 0.26 nm determined by X-ray diffraction analysis for a *p*-mercaptobenzoic acid-capped GNP.^33^ The average distributions in distance between AU and sulfur atoms in the equilibrated systems (Fig. S2 in the Supporting Information) show, taking into account the uneven GNP surface, that after equilibration the sulfur atoms remain at the expected distance from the nominal surface. Moreover, we note that, because of the polarizable gold atoms, not only the deprotonated thiol group, but also the charged amino and carboxylate functional peptide groups interact with the GNP surface, but with somewhat weaker van der Waals interaction energies than the thiol group. Indeed, the amino groups are found at similar distance to the GNP surface as the sulfur atoms (Fig. S2). This is in agreement with the experimental observation that protonated amino groups interact with gold surfaces^34^ and contribute to anchoring the peptides on the GNP.^9^

Taking into account the experimentally determined peptide capping density to be modelled, a number of peptides (for symmetry reasons limited to values of 4n^2^+2, where n is an integer) was roughly equidistantly placed around a gold nanoparticle of desired size.

To investigate the effect of the capping density on the structure of a CALNN monolayer on a GNP, molecular dynamics (MD) simulations for CALNN-capped GNPs at high and low capping density were performed. CALNN peptides were arranged around a spherical GNP of diameter 10 nm at a capping density of either 2.5 (N_peptide_ = 786) or 1.8 (N_peptide_ = 578) peptides/nm^2^. These structures were placed in a cubic box with sides of length 17 nm; 786 or 578 sodium ions were added to neutralize the system. Water molecules were placed around the CALNN-capped gold nanoparticle to obtain a water density of ~1 g/cm^3^ after full equilibration. A schematic representation of the starting structures of the two systems is shown in Figure 1.

**Figure 1.**
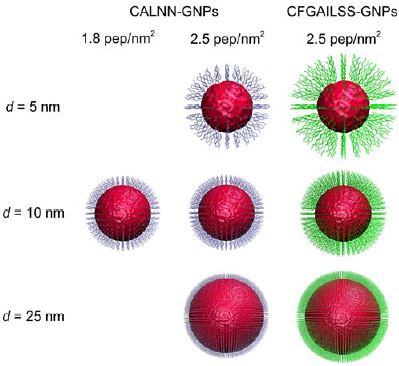
Snapshots of CALNN-and CFGAILSS-capped gold nanoparticle starting structures; (left) CALNN-capped 10 nm GNP at a capping density of ~1.8 peptides/nm^2^; (center) CALNN-capped 5,10 and 25 nm GNPs at a capping density of ~2.5 peptides/nm^2^; (right) CFGAILSS-capped 5, 10 and 25 nm GNPs at a capping density of ~2.5 peptides/nm^2^. For clarity, water and sodium ions are not shown and the display scales are adjusted for each particle size.

To study the effect of GNP curvature on the structure of CALNN and CFGAILSS monolayers on gold nanoparticles, MD simulations for CALNN-and CFGAILSS-capped GNPs of 5, 10 and 25 nm in diameter were carried out. The peptide capping density was kept as close as possible to the experimental values^27,35^ of 2.4 and 2.55 peptides/nm^2^ for CALNN-and CFGAILSS-capped GNPs, respectively (N_peptide_ = 198, 786, and 4626 (CALNN) or 4902 (CFGAILSS) for the three nanoparticle sizes). These structures were placed in a cubic box with sides of length 12, 17 or 32 nm, respectively. Similarly to the previous systems, sodium ions were added to neutralize the system and the water density was ~1 g/cm^3^ after full equilibration. It is worth remarking that the size of the cubic box was chosen in order for all the systems to have roughly the same distance between the fully extended peptide and the walls of the box. A schematic representation of the starting structures of the six systems is shown in Figure 1.

The peptide-capped GNP was first energy-minimized in vacuum with the steepest descendent method. Consecutive NVT ensembles were then applied using the Berendsen method^36^ to couple the temperature at 298 K; the gold nanoparticle and the peptides were coupled separately. The Verlet cut-off scheme at 1.2 and 1.4 nm for short and long-range neighbor lists was used and the lists were updated every 100 fs. Both Coulomb and van der Waals interactions were calculated with a cut-off at 1.2 nm. The peptide-capped GNP was then solvated using the TIP3P water model^37^ at a density of ~1 g/cm^3^ and the negative charges of the carboxylate functional groups neutralized by addition of sodium ions. Equilibration of the whole system in an NVT ensemble was carried out using the Berendsen method^36^ to maintain the temperature at 298 K and coupling the gold nanoparticle, the peptides, and the solvent together with the ions in three separated groups. The short-range and long-range neighbor lists had a cut-off at 1.2 and 1.4 nm using the group cut-off scheme and were updated every 20 fs. Long-range electrostatic interactions were treated with the Particle-mesh Ewald method^38^ with a Fourier grid spacing of 0.12 nm. The short-range van der Waals interactions were calculated using a switching function between 1.0 and 1.2 nm. The bond lengths were constrained via the LINCS algorithm,^39^ a time step of 1 fs was used and periodic boundary conditions were adopted. Depending on the system size, equilibrium was reached after different simulation times, as assessed by macroscopic parameters, such as the total energy, being constant over time. Hence, structures with 5 nm GNPs required at least 20 ns to equilibrate, whereas those with 10 or 25 nm GNPs took more than 30 or 40 ns, respectively; data were then collected for the subsequent 10 ns (“production run”). Furthermore, in order to test the reproducibility of the computational results, the CFGAILSS-capped 10 nm GNP was also simulated with different starting conditions, i.e., using the Berendsen method^36^ to couple the temperature at 348 K for the first set of NVT ensembles applied on the gold nanoparticle and the peptides. All the analyses were done using programs in the GROMACS package and custom-written software. For secondary structure assignment, the DSSP algorithm^40^ was employed. Visual Molecular Dynamics version 1.9.1^41^ was used to visualize and render the structures and create the movie available in the SI.

## RESULTS AND DISCUSSION

### Effect of Capping Density on the Structure of a CALNN Monolayer (Experimental Investigation)

The CALNN pentapeptide was designed to form a self-assembled monolayer around GNPs and thus impart colloidal stability.^9^ Amino-acid analysis revealed that the peptide capping density, hence the monolayer compactness, is dependent on the concentration of peptide used during the ligand exchange procedure.^35^ We previously investigated the secondary structure of CALNN monolayer by FTIR spectroscopy and reported both a random coil^27^ and a straight conformation.^42^ In the latter work, we hypothesized a difference in the peptide capping density to underlie these divergent observations. Here, a more systematic experimental study on CALNN-capped GNPs at different capping densities is presented in parallel with molecular dynamics simulations.

FTIR spectroscopy was employed, first, to estimate the CALNN capping density and, second, to characterize the secondary structure of the monolayer. In particular, the amide I’ region (1600 to 1700 cm−^1^), mainly associated with the backbone carbonyl stretching vibration, which gives information on the backbone conformation,^43^ was analyzed. The amide I’ band area was used to estimate the peptide capping density (Supporting Information for details and unscaled FTIR spectra, Figure S3), taking into account the GNP concentration, as determined by UV-vis spectroscopy, and the IR surface selection rules, which quantitatively describe the strength of the IR-active modes of small molecules bound to a metal surface or nanoparticle, as described in detail previously.^42^ The estimated capping density for CALNN-capped GNPs at high and low capping density are 2.1 and 1.4 peptides/nm^2^, in reasonable agreement with those reported previously, using amino-acid analysis, for CALNN-GNPs prepared with the same ligand exchange procedures (2.4 and 1.7 peptides/nm^2^, respectively).^35^ The FTIR spectra of CALNN peptide in solution and of CALNN-capped GNPs at high and low capping density are shown in Figure 2. The amide I’ band of CALNN peptide at 1651 cm^−1^, corresponding to a random coil conformation,^44^ shifts to 1648 cm^−1^ and 1641 cm^−1^ when constituting the low and high capping density monolayer, respectively. The position of the former is still indicative of a largely disordered peptide monolayer, whereas the latter is denoting a straight conformation, either polyproline II (PPII) or β-strand;^45–47^ although one of the bands observed for parallel β-sheets is at a very similar position,^27^ the accompanying high-frequency band is missing here, allowing us to rule out this structure. The PPII motif^48^ is not only found in proline-rich polypeptides, but also in nominally disordered polypeptides^45,49^ and is characterized by the lack of internal hydrogen bonding. As for the β-strand, it corresponds to a single peptide strand in a β-conformation, i.e., a quite extended structure lacking the stabilizing cross-strand hydrogen bonds that would form a β-sheet. In the absence of an internal network of stabilizing hydrogen bonds, both motifs are thought to be stabilized by hydrogen bonds between the peptide backbone and the surrounding water molecules^50^ and the amino acids within such structures expose to the solvent 60% more polar surface area compared to other residues.^51^ Hence, the amide I’ band in the IR spectrum of CALNN-capped GNPs with low capping density indicates a capping layer consisting of largely disordered peptides, whereas that of high capping density CALNN-capped GNPs, while indicating a straight conformation, is not in agreement with a β-sheet-like structure; however, the experimental results cannot distinguish between PPII and β-strand conformations for this system.

**Figure 2.**
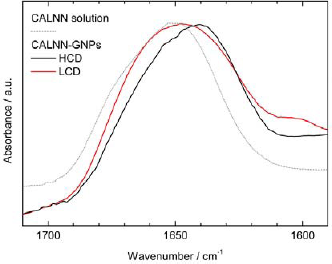
FTIR spectra in the amide I’ region of CALNN peptide in solution and of 11 nm CALNN-capped gold nanoparticles at high and low capping density (HCD and LCD, respectively). All spectra have been scaled to the same maximum absorbance.

### Effect of Capping Density on the Structure of a CALNN Monolayer (Computational Investigation)

Molecular dynamics simulations of two systems that model the high and low capping density CALNN-capped GNPs were performed. The former and the latter had a CALNN capping density of 2.5 and 1.8 peptides/nm^2^ on a GNP of 10 nm diameter, respectively (see Computational Model section), to roughly reflect the ~30% difference in capping density of the CALNN-capped GNPs analyzed by FTIR spectroscopy. The starting structures of the two simulated systems are shown in Figure 1.

To establish the validity of the computational model, the secondary structure of the simulated high and low capping density CALNN monolayers on GNPs was compared to the IR experimental findings. The time-evolving secondary structure of CALNN peptides within the high and low capping density monolayers, based on DSSP assignment,^40^ is shown in the top panels of Figure 3 (see Supporting Information, section S4, for comments on the DSSP algorithm). It is worth noting that the secondary structure is stable over time in both simulated systems (Figures 3A,B). The overall content of well-defined structures, as identified by the DSSP algorithm, is very small in both systems, with more than 90% of the content identified as “random coil”. On the other hand, there is a noticeable difference between the low and high capping density monolayers - whereas for the former less than 2% of the residues are found in β-structures (i.e., β-sheet or β-bridge), slightly more (5%) are in these conformations for the latter. Thus, the high capping density monolayer appears to be more structured. However, as discussed above, the amide I’ band in the FTIR spectrum of high capping density CALNN-capped GNPs is indicative of a PPII or β-strand conformation, both extended secondary structure motifs lacking intermolecular hydrogen bonding. Because the DSSP secondary structure assignment^40^ is mostly based on the recognition of hydrogen bonds and therefore fails in the identification of such motifs, the backbone ϕ and ψ torsion angles for the two simulated systems were also analyzed to gain more detailed information on the local backbone conformation (Figure 3, bottom; see also section S5 in the Supporting Information for details of the residues considered and a quantitative analysis of the Ramachandran plots). Characteristic ϕ and ψ ranges are associated with well-defined secondary structure motifs.^52^ For instance, typical values for antiparallel β-sheets are ϕ = −147°, ψ = +145°,^53^ for parallel β-sheets ϕ = −116°, ψ = +112°,^53^ for PPII ϕ = −75°, ψ = +145° and for right-handed α*-* helices ϕ = −60°, ψ = −45°. Thus, extended backbone structures are found near the top left corner of the Ramachandran plot. Figures 3 C and D show that a significant fraction of residues are found in this region for both systems. However, there is a significant shift of population from the helical region to the region of extended conformations, indicating a “straightening” of the peptides, upon increasing the capping density. It should be noted that Ramachandran plots show the conformation of individual residues and population in the α-helical region does not indicate formation of helical secondary structures, which does not occur for CALNN, see Figure 3A,B. Thus, the structure information arising from DSSP and Ramachandran analyses are complementary and indicate a more structured monolayer with more extended peptides in the case of GNPs with a higher capping density of CALNN peptide; most of these extended peptides, however, do not form good agreement with the conclusions drawn from the experimental FTIR spectra. Moreover, the Ramachandran plot suggests that the extended β-strand conformation is somewhat preferred over the PPII conformation in high capping density CALNN monolayers, although both conformations are found.

**Figure 3.**
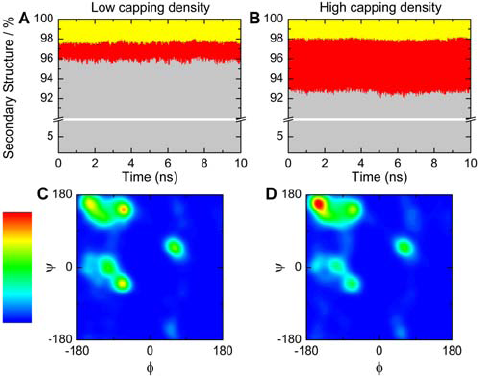
Secondary structure of CALNN monolayers, based on (top) DSSP assignment over time and (bottom) Ramachandran analysis of the final structure for 10 nm CALNN-capped GNPs at low (A,C) and high (B,D) capping density. Secondary structure color code in A and B: grey, random coil; red, β-structures (i.e., β-sheet or β-bridge); yellow, turn. The Ramachandran plots are colored by occupancy; the intensity scale range is from 0 (dark blue) to 0.00012 (red) residue probability/angle^2^.

The fact that the simulated secondary structure content well reflects the IR experimental findings supports the validity of the computational model, from which further insights into the monolayers can be gained.

To estimate how extended the CALNN peptides are, the distance distribution between the N− and C− peptide terminal atoms was examined (Figure 4A). For both systems, a similar fraction of peptides was found to be significantly folded back onto themselves (d_N−C_ ≤ ~1 nm). On the other hand, in the region of more extended conformations (d_N−C_ ≥ ~1 nm), there are clear differences; for the high capping density system, the d_N−C_ distribution is narrower and shifted towards the straight conformation (d_N−C_ ~1.6 nm for an antiparallel β-strand^53^ or ~1.5 nm for a PPII structure^54^ consisting of 5 residues), reflecting the need for more ordered conformations in a monolayer of higher compactness. These findings are supported by visual inspection of the structures, see the snapshots in Figure 4, which show that both high (Figure 4B) and low (Figure 4D) capping density monolayers are characterized not only by some back-folding of peptide chains, but also by peptides in an extended conformation, although the latter are more dominant in the high capping density system.

**Figure 4.**
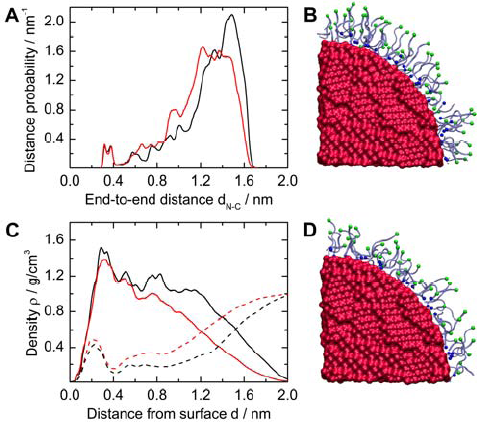
Extension of CALNN peptides analyzed by (A) distribution of the distance d_N−C_ between the peptide N-and C-terminal atoms and (C) radial density of peptide (solid lines) and water (dashed lines) atoms at distance d from the surface of the GNP. Black and red curves refer to the high and low capping density system, respectively. All results were averaged over 10 ns simulation. Snapshots of final structures of 10 nm CALNN-capped GNPs at high and low capping density are illustrated in B and D, respectively (shown is one-eighth of the structure); the peptide N-and C-terminal atoms are rendered in blue and green, respectively.

Further information about the internal structure of the monolayers can be gained from the radial density profiles of peptide and water atoms shown in Figure 4C. For the high capping density system, the density profile of peptide atoms (Figure 4C, solid black line) shows an almost constant value of ~1.2 g/cm^3^ up to ~1.2 nm from the GNP surface, which is close to the widely accepted density for polypeptides^55^ of 1.4 g/cm^3^; the radial density profile of water atoms (Figure 4C, dashed black line) indeed shows that not much solvent is present in this region, confirming the presence of a compact peptide monolayer. The gradual decay of the peptide atom density at larger distances from the gold surface reflects the mobility and disorder of the peptide carboxyl ends. For the low capping density system (Figure 4C, red lines), on the other hand, no such region of constant peptide atom density was found; the decay is more gradual and is accompanied by a more gradual increase of the water content in the capping layer, confirming the significantly more disordered nature of the low capping density monolayer, in spite of the fact that the capping density is reduced by only 26%. It is interesting to note, however, that almost the same amount of water appears to be trapped near the GNP surface for both monolayers.

The radial density profiles also allow the determination of an effective capping layer thickness, which is a quantity that could be compared to experimental results, obtained e.g., by Dynamic Light Scattering or Differential Centrifugal Sedimentation (DCS) techniques. For the sake of avoiding ambiguity, we here define the effective thickness as the distance from the GNP surface at which the peptide atoms have the same density as the solvent, which yields values of 1.21 and 1.52 nm for high and low capping density CALNN-capped GNPs, respectively. Interestingly, this estimated thickness of the high capping density system is only slightly less than the predicted length of 1.7 nm for an antiparallel β-strand consisting of 5 residues,^53^ and in very good agreement with the predicted length of a 5-residue PPII structure.^54^

Thus, FTIR spectroscopy showed that even a relatively small change of the CALNN capping density does have a significant effect on the structure of the monolayer on a GNP, suggesting that CALNN peptides have to adopt a more extended structure in order to organize in a monolayer with high capping density. The computational model proposed here does not only fully reproduce this effect, but also allows more detailed insight into the secondary structure motifs and structural properties of CALNN monolayers at different capping densities. This provides molecular details which have not yet been observed experimentally, including a pocket of water close to the gold surface and a small proportion of peptides in a loop configuration with their C-terminus in contact with the gold surface.

### Effect of Surface Curvature on the Structure of CALNN and CFGAILSS Monolayers

In previous experimental work,^27^ we investigated the secondary structure of CALNN and CFGAILSS peptide monolayers on GNPs of different sizes, i.e., 5, 10 and 25 nm in diameter, by means of FTIR and solid-state nuclear magnetic resonance (ssNMR) spectroscopies. CALNN was found to adopt a random coil conformation independent of the GNP size, whereas CFGAILSS, which forms extended amyloid fibers with antiparallel β-sheet conformation in solution, showed a propensity to form parallel β-sheets when attached to GNPs which depended on GNP size, i.e., curvature, with almost no β-sheets observed on 5 nm GNPs, some β-sheet formation on 10 nm GNPs, and significant β-sheet propensity on the 25 nm large nanoparticles. A geometrical model was proposed in order to rationalize the role of the nanoparticle curvature on the secondary structure of CFGAILSS. According to this model, the number of adjacent peptides that can form hydrogen bonds and therefore be involved in parallel β-sheets is dictated by the nanoparticle curvature. Thus, more extended parallel β-sheets are observed on the larger nanoparticles, presenting the lower curvature. In the IR investigation of Mandal et al„ a Leu-rich peptide was found to be more prone to form α-helices on gold nanoparticles presenting lower curvature, possibly because of a greater interpeptide distance.^56^ Therefore, these IR studies highlighted the importance of the peptide sequence and nanoparticle size when designing peptide-capped gold nanoparticles with well-defined secondary structure motifs.

Here, the experimentally characterized CALNN-and CFGAILSS-capped GNPs with 5, 10 and 25 nm diameter were translated into six computational systems illustrated in Figure 1. The peptide capping density reflected as closely as possible the experimental values^27–35^ measured by amino acid analysis of 2.4 and 2.55 peptides/nm^2^ for CALNN-and CFGAILSS-capped GNPs, respectively (see Computational Model section).

The peptide secondary structures found in the computational model can be directly compared to the experimental results. Figure 5 illustrates that the secondary structure remains constant over the 10 ns “production run” in all systems and that significant differences between GNPs of different sizes can be observed for CFGAILSS monolayers. For instance, with the increase of the GNP diameter from 5 to 10 nm (Figure 5D,E), the content in β-structures in CFGAILSS monolayers sees a moderate growth from ~5.5 to 11%, whereas a more dramatic increase to ~21% can be observed upon increasing the GNP size to 25 nm (Figure 5E,F). This increase of β-structures is accompanied by a loss of bending conformations. For CALNN monolayers, the changes in secondary structure motifs with an increase in GNP size are much smaller (Figure 5A-C). Thus, based on DSSP secondary structure assignment, the simulations are in remarkably good agreement with the FTIR and ssNMR observations of ref.^27^. In the case of CFGAILSS systems, they confirm a small difference in the amount of β-structures between 5 and 10 nm GNPs and that the highest β-content is on the GNP with lower curvature, i.e., the 25 nm GNP. In the case of CALNN monolayers, they indicate a small increase in β-structures on the larger GNPs, but confirm a mostly random coil conformation, independent of the nanoparticle size. However, as previously mentioned, the DSSP secondary structure assignment is mostly based on the recognition of hydrogen bonds between amino acids,^40^ and thus cannot account for secondary structure motifs that are not denoted by intermolecular hydrogen bonding, such as PPII and β-strand structures.

**Figure 5.**
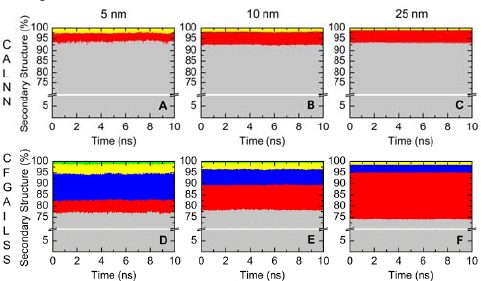
Time-evolving secondary structure for CALNN (top) and CFGAILSS (bottom) monolayers on 5 (A,D), 10 (B,E) and 25 nm (C,F) GNPs, based on DSSP assignment. Secondary structure color code: grey, random coil; red, β-structures (i.e., β-sheet or β-bridge); blue, bend; yellow, turn; green, helix.

To derive conclusions on these structures, the backbone ϕ and ψ torsion angles were analyzed. With an increase of the GNP diameter from 5 to 10 nm, the Ramachandran plots of CALNN systems (Figure 6A,C, see also Table S1 in the Supporting Information for a quantitative analysis of the Ramachandran plots) indicate a shift of population from the PPII (ϕ= −75°, ψ = +145°) and particularly the right-handed α-helical (ϕ = −60°, ψ = −45°) regions to the antiparallel β-sheet region (ϕ= −147°, ψ = +145^0^), indicating a “straightening” of the peptides. Thus, decreasing the curvature forces the peptides into a more extended conformation, similar to the effect of increasing the capping density on 10 nm GNPs, as discussed above. On the other hand, almost no difference is observed between 10 and 25 nm CALNN-capped GNPs (Figure 6C,E), in agreement with the DSSP analysis results of Figure 5. In the case of the CFGAILSS systems, a similar, but much more pronounced shift of population from the right-handed α-helical to the extended region, and within the extended region a shift from the PPII to the antiparallel and parallel β-sheet regions is observed with an increase of the GNP size, (Figure 6B,D,F); it is particularly noteworthy that the increase from 10 to 25 nm GNP diameter leads not only to a significant increase of population in the β-sheet region in general, but to a pronounced redistribution of the β-sheet conformations towards the parallel β-sheet region (ϕ = −116°, ψ = +112°). Thus, the structural information arising from DSSP and Ramachandran analyses support each other, and show that the simulations are in good agreement with the conclusions which had been drawn from FTIR and ssNMR spectroscopy.

**Figure 6.**
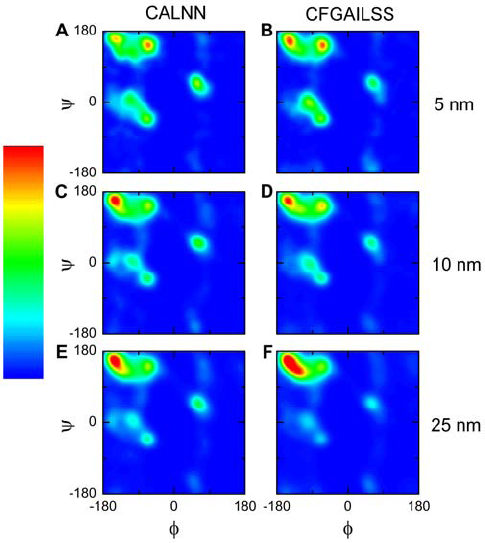
Ramachandran plots of the final structure for CALNN (left) and CFGAILSS (right) monolayers on 5 (A,B), 10 (C,D) and 25 nm (E,F) GNPs. The plots are colored by occupancy; the intensity scale range is from 0 (dark blue) to 0.00012 (red) residue probability/angle^2^.

The fact that the simulated secondary structure content well reflects the experimental findings supports the validity of the computational model, from which further insights into the monolayers can be gained.

To assess the structure at the peptide level, the distance distribution between the N-and C-peptide terminal atoms was examined. The profiles for CALNN-capped GNPs (Figure 7A) highlight that a portion of CALNN peptides is back-folding on the GNP surface (d_N−C_ ≤ ~1 nm). Also, they confirm, in line with the results for the secondary structure, that with an increase of GNP size, hence a decrease of curvature, there is some degree of “straightening” - with increasing GNP size, the d_N−C_ distribution narrows and shifts towards the straight conformation (d_N−C_ ~1.6 nm for an antiparallel β-strand^53^ or ~1.5 nm for the PPII structure^54^ consisting of 5 residues). This curvature-induced straightening is comparable to the effect of increasing the capping density, see above; here, the reduced curvature of the larger GNPs pushes the peptide C-termini together, which leads to a more compact capping layer and thus requires a more ordered peptide conformation. It is noteworthy that most of this effect occurs upon an increase of the GNP diameter from 5 to 10 nm, whereas a further increase has almost no effect on the distribution of d_N−C_. In contrast, CFGAILSS-capped GNPs (Figure 7B) show much less peptide back-folding, with almost no population in the region d_N−C_ ≤ ~1.5 nm, and a much more pronounced shift to almost fully extended peptide conformations (d_N−C_ = 2.7 nm for an antiparallel β-strand consisting of 8 residues)^53^ with increasing GNP size; unlike CALNN, the effect is more pronounced when comparing 10 and 25 nm GNPs. This again is in full agreement with the secondary structural analysis of the simulations, as well as with the experimental results for a CFGAILSS capping layer on GNPs of increasing size. Thus, in addition to the higher compactness of the capping layer as a consequence of decreased GNP surface curvature, which is observed even for an unstructured peptide like CALNN, there is another effect operating for CFGAILSS: this peptide has a significant propensity for forming β-sheets, but when the N-termini are attached to a high curvature surface, the backbones of adjacent peptides are too far shifted relative to each other to allow inter-strand hydrogen bond formation. Increasing the GNP size, and hence decreasing the surface curvature, reduces this misalignment and thus leads to increased β-sheet formation which provides an additional “driving force” for peptide straightening. Although some β-sheet formation already can occur on 10 nm GNPs, the surface curvature still prevents formation of extended β-sheets; only on the 25 nm GNPs can this be observed.

**Figure 7.**
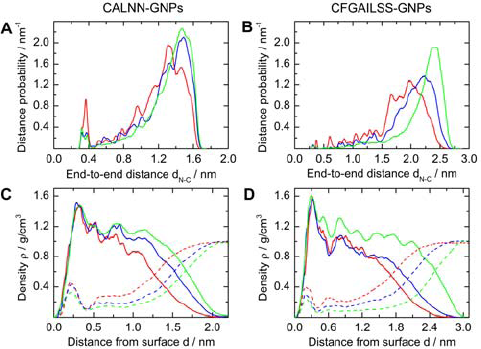
Extension of CALNN (left) and CFGAILSS (right) peptides analyzed by (A,B) distribution of the distance d_N−C_ between the peptide N-and C-terminal atoms and (C,D) radial density of peptide (solid lines) and water (dashed lines) atoms at distance d from the surface of the GNP for CALNN-and CFGAILSS-capped 5 (red lines), 10 (blue lines) and 25 nm (green lines) GNPs, averaged over 10 ns simulation.

To gain more information about the structure of the monolayer and to estimate its thickness, the radial density of peptide and water atoms was analyzed (Figures 7C,D). As discussed before, CALNN peptides in a high capping density monolayer on a 10 nm GNP display a compact monolayer characterized by a peptide density which is close to the widely accepted density for polypeptides^55^ of 1.4 g/cm^3^ and low water content. Almost the same density profile is found for the larger 25 nm GNPs, whereas for the small (5 nm) GNPs, a less compact peptide layer, characterized by a more gradual decay of the peptide density and higher water content, was found, which is reminiscent of the low capping density CALNN layer on 10 nm GNPs. The effective thickness, i.e., the distance from the GNP surface at which the peptide and water density are equal, increases from 1.25 nm for the 5 nm GNPs to 1.52 nm (10 nm GNPs) and 1.67 nm (25 nm GNPs). On the other hand, the effect of nanoparticle curvature on the compactness of a CFGAILSS monolayer, as indicated by the radial density, is much more pronounced than for a CALNN monolayer. While the two monolayers show a similar gradual decrease of peptide density on 5 nm GNPs, on 10 nm GNPs the CFGAILSS monolayer is less compact, with a significantly lower peptide density, than a CALNN monolayer with the same capping density. Only for 25 nm GNPs does the CFGAILSS capping layer become as compact and dense as the CALNN monolayer, with a high peptide density close to that of polypeptides and low water content up to ~2 nm from the GNP surface. The effective CFGAILSS layer thickness increases from 1.71 nm for the 5 nm GNPs to 2.01 nm (10 nm GNPs) and 2.50 nm (25 nm GNPs). The latter value is in good agreement with the length of a parallel β-sheet consisting of 8 residues (2.60 nm).^53^ As already noted above for CALNN monolayers of varying packing density, some water appears to be trapped near the GNP surface also for CFGAILSS monolayers, with some decrease of the water density near the surface as the GNP curvature decreases.

### Insights into the β-structures within CFGAILSS Monolayers

Since, as discussed above, most of the straightening of CALNN peptide is not accompanied by inter-strand hydrogen bonding and hence is not well captured by DSSP analysis, we focused on CFGAILSS monolayers to gain more detailed insights into secondary structure motifs. Figure 8 shows the percentage of secondary structure, as identified by DSSP analysis,^40^ for each residue in CFGAILSS peptide chain (due to DSSP analysis limitations no secondary structures are assigned to the first and the final two residues, see Supporting Information, section S4, for more comments on the DSSP analysis). The profiles illustrate that decreasing the GNP curvature leads to an increase of β-structures and decrease of bending predominantly for the inner segment Phe-Ile, with Gly-Ala having the largest β-content. Moreover, the average length of β-structures is 1.3,1.8 and 2.3 residues for 5,10 and 25 nm CFGAILSS-capped GNPs, respectively. Hence, more extended segments of peptide participate in β-structure formation on the nanoparticles with lower curvature, i.e., larger size.

**Figure 8.**
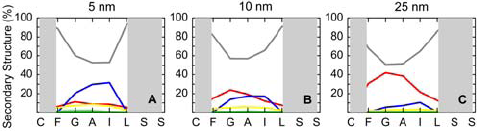
Secondary structure content per amino acid for CFGAILSS monolayer on 5 (A), 10 (B) and 25 nm (C) GNPs, based on DSSP assignment, averaged over the last 2 ns of the simulation. The grey areas depict the amino acids not identifiable with secondary structure motifs due to limitations of the DSSP algorithm. Secondary structure color code: grey, random coil; red, β-structures (i.e., β-sheet or β-bridge); blue, bend; yellow, turn; green, helix.

The snapshots in Figure 9 show that not only does the number of parallel β-sheets increase with a decrease in nanoparticle curvature, but they also appear to associate into more extended parallel β-sheet domains.

**Figure 9.**
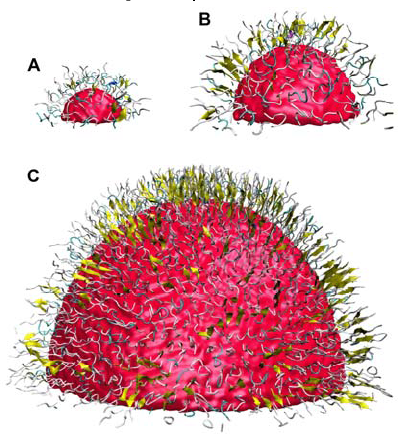
Snapshots of a portion of the final structures of CFGAILSS-capped 5 (A), 10 (B) and 25 (C) nm GNPs. Peptides are rendered with the VMD “New Cartoon” drawing method and colored per secondary structure motif; β-sheets are illustrated as yellow arrows. The images are to scale and shown in perspective.

To explore the existence of such supra-molecular domains within CFGAILSS monolayers, the distribution of β-structure probability is displayed on polar charts (Figure 10; polar charts of the other hemispheres are shown in the Supporting Information, Figure S5). The black dots indicate the position of the backbone center of mass for each peptide. The local distributions of β-structure probability clearly illustrate that a high nanoparticle curvature (Panel A) limits the size of β-structure domains to not more than two or three peptides, since the curvature leads to misalignment of neighboring peptide backbones, whereas a low nanoparticle curvature allows the formation of extended β-structure domains (Panel C). Also, the β-structures appear to be randomly distributed across the nanoparticles. It is worth noting that the spatial distribution of β-structures does not change much during the 10 ns “production run”, indicating that the full conformational landscape is not accessed over the ns-timescale of molecular dynamics simulations, in agreement with the known time scales of secondary structure equilibration dynamics, see below. The results for a CFGAILSS-capped 10 nm GNP simulated with different starting conditions (see Computational Model section for details), show indeed a similar β-structure content as the 10 nm CFGAILSS-capped GNP previously discussed, i.e., 9 and 11%, respectively, but a different spatial distribution of β-structures, see Figure S7 in the Supporting Information. This structure represents another one of the many local minima on the shallow conformational landscape with different local structures, but very similar average properties, which make up the equilibrium ensemble. Because of the time scale of secondary structure equilibration dynamics, see below, transitions between these local minima occur on longer time scales than those available to molecular dynamics simulations. However, the observation that the average properties of the two structures simulated with different starting conditions are highly similar confirms that the results obtained here are valid and relevant for comparison with experimental results, which necessarily are ensemble averages. For reference, the polar charts for the CALNN monolayers are reported in Figure S6; these show significantly lower β-structure probabilities, in agreement with the results presented above, and consequently no extended β-structure domains.

**Figure 10.**
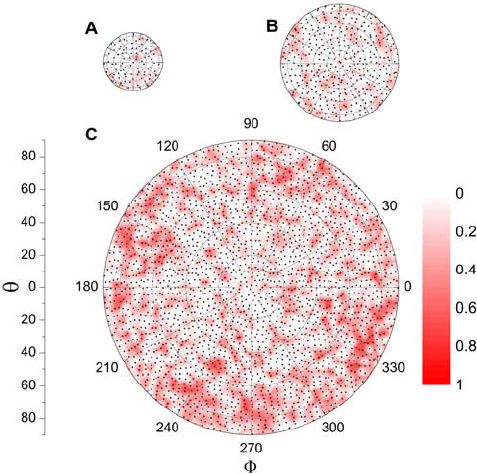
Spatial distribution of β-structure probability, as assessed by DSSP analysis, displayed on polar charts, where Φ and θ indicate the azimuthal and polar angle in spherical coordinates, respectively, for the final structures of CFGAILSS-capped 5 (A), 10 (B) and 25 nm (C) GNPs. The black points indicate the polar position of the backbone center of mass of each peptide; the color contour shows the local β-structure probability.

Close inspection of the distribution of peptides in the domains of high β-structure content within the CFGAILSS capping layer on 25 nm GNPs suggests that these peptides arrange in regular repeat patterns in two dimensions (Figure 11B). Many of these peptides are found in lines with a typical inter-peptide distance of ~0.5 nm, which is in good agreement with the distance between two β-strands in a parallel β-sheet, i.e., 0.47 nm. Furthermore, these lines of peptides often run parallel to each other, with a distance on the order of ~1 nm, corresponding to the inter-β-sheet distance in amyloid fibrils, which consist of stacked β-sheets.^57^ Thus, we suggest that on 25 nm GNPs CFGAILSS peptides arrange in an amyloid-like structure consisting of stacked parallel β-sheets. The stacking of parallel β-sheets is in contrast to most amyloid structures, including those formed by CFGAILSS peptides in solution,^27^ which consist of antiparallel β-sheets; however, binding of the peptides to the GNP surface by their N-terminus necessarily prevents the formation of antiparallel β-sheets. For a more quantitative confirmation of this suggestion, the radial distribution function of the backbone-backbone distances was analyzed (Figure 11A). The nearest neighbor distribution for peptides with no β-structures shows a maximum at ~0.78 nm; interestingly, this corresponds to a capping density of less than 2 peptides/nm^2^, which is much smaller than the average capping density (2.5 peptides/nm^2^), showing that formation of β-sheets occurs in domains of increased capping density, which leads to depletion of the remaining surface. This suggests that movement of the sulfur atoms on the gold surface may be critical to the formation of secondary structure motifs. Simulations in which sulfur atoms were strongly bound to the gold surface were also performed (for details of these simulations, see Supporting Information, section S2). In contradiction to the experimental findings^27^ and the results obtained using the GolP-CHARMM force field, CFGAILSS-capped 10 nm GNP simulated with this computational model showed no significant secondary structure formation (Figure S8), thus confirming the importance of the mobility of peptides on the surface (from the starting to the final structure, the sulfur atoms moved on average by ~0.1 nm in the CHARMM-METAL model and by ~0.4 nm in the GolP-CHARMM model).

**Figure 11.**
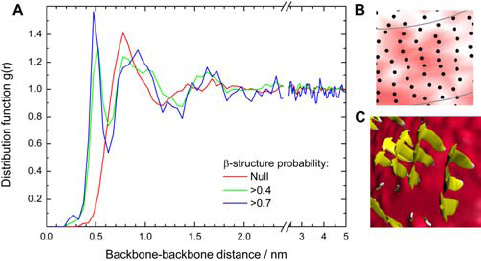
(A) Radial distribution functions of the backbone-backbone distances for peptides with different β-structure probabilities in a CFGAILSS-capped 25 nm GNP; the profiles are the result of averaging over six equilibrated structures; (B) example of a β-structure domain denoted by an amyloid-like arrangement of peptides (zoom-in of Fig. 10C); (C) topview snapshot of parallel β-sheets on gold surface, showing amyloid-like structure.

In contrast, the radial distribution functions for peptides with high β-content (>0.7) are characterized by maxima at 0.5 and 0.95 nm, corresponding to the inter-β-strand and inter-β-sheet distances found in amyloid fibrils, which confirms the conclusion drawn above from a visual inspection of the polar plot. Figure 11C shows a snapshot of such parallel β-sheets on the nanoparticle surface that illustrates the amyloid-like arrangement of peptides within some β-structures domains. For reference, the radial distribution function of the backbone-backbone distances for the CALNN-capped 25 nm GNP is reported in Figure S9; it shows a similar distribution for peptides with no β-structures as found for the CFGAILSS-capped 25 nm GNP, whereas no maximum at ~1 nm is found even for peptides with the highest β-sheet content, in agreement with the absence of amyloid-like structures concluded on by visual inspection of the structure.

### Peptide Dynamics

The Supporting Information contains a movie of the CFGAILSS-capped 10 nm GNP, in which the secondary structure of CFGAILSS monolayer is visualized every ps over the last 100 ps of the 10 ns “production run” (the peptides are rendered with the VMD “New Cartoon” drawing method and colored per secondary structure motif; β-sheets are illustrated as yellow arrows and, for clarity, water and sodium ions are not shown). This clearly shows that the β-structures and in particular the supra-molecular domains into which they associate do not change much over time; this is valid for the full 10 ns “production run”, as evidenced by the spatial distributions of β-structure probability at different times shown in Figure S10 in the Supporting Information. Moreover, as previously discussed, the CFGAILSS monolayer on a 10 nm GNP simulated with different starting conditions revealed a similar average content of β-structures, but with a completely different spatial distribution (Figure S7), which again was found to be largely constant on the 10 ns time scale. Thus, it can be concluded that over the ns-timescale which is available to molecular dynamics simulations the individual peptides do not probe the full conformational landscape which they would sample over longer time scales. Instead, the system remains within a local minimum of the conformational landscape, corresponding to a particular distribution of β-structures. This is not surprising, since it is well known from experimental evidence that equilibration of secondary structures is slower than 10 ns. Thus, the folding/unfolding dynamics of α-helices occur on the 100 ns time scale^45^ and those of β-sheet structures are even slower.^58,59^ Hence, the simulations reported in this study can be considered as equilibrated with respect to the overall properties (ensemble averages), but are not ergodic, i.e., the localization of CFGAILSS β-structures averaged over the production run is not representative of an ensemble average (which would be homogenous over the surface due to the spherical symmetry). For a further analysis, the time autocorrelation of the β-structure probability for each of the five residues in the CFGAILSS peptides which can be analysed by DSPP was calculated and is shown in Figure 12.

**Figure 12.**
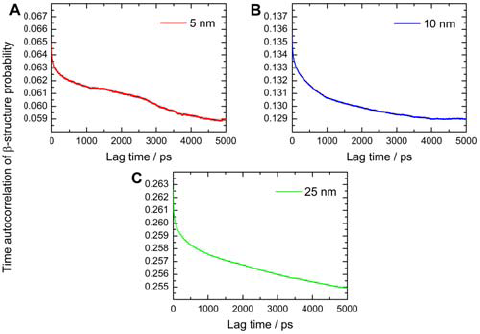
Time autocorrelation of β-structure probability resulting from the average of the time autocorrelation functions for F,G,A,I,L residues, averaged over all peptides, for CFGAILSS monolayer on 5 (A), 10 (B) and 25 nm (C) GNPs; β-structure probability extracted from DSSP analysis in time steps of 1 ps over the 10 ns production run.

Fig. 12 shows that only the onset of the autocorrelation decay is captured. Hence, since there is no reason to assume any particular functional time dependence for the autocorrelation functions, no exact quantitative analysis is possible, such as the determination of an autocorrelation time. However, analysis of the onset allows the prediction that autocorrelation times exceed several 100 ns, in full agreement with the expectations for β-sheet (un)folding.^58,59^

## CONCLUSIONS

Here, we have presented a computational model for peptide-capped gold nanoparticles that account for the motion of sulfur atoms over the nanoparticle surface and dynamic polarization of gold atoms. We have pointed out the crucial importance of the cysteine sulfur atoms’ mobility for the formation of secondary structure motifs and described the set-up and validation of the computational model using experimentally well-characterized systems as benchmarks. Furthermore, we have shown that MD simulations provide insights not only into the peptides’ structural and dynamic properties, but also into the distribution of secondary structure domains on the gold surface and into the peptides’ arrangement within such domains. Therefore, while elucidating the structure of peptide monolayers on gold nanoparticles is experimentally challenging and the detailed insights attainable often limited, our study shows that a combination of experiments and modelling is quite compelling for determining the monolayer structure at different molecular levels. In the future, the computational model proposed here will be used to predict the structure of other peptide monolayers on gold nanoparticles and help with the experimental design of peptide-capped gold nanoparticles with well-defined structure motifs, which could also be assembled into objects with complexity similar to the one found in proteins.

## MATERIALS AND METHODS

### Gold Nanoparticles

Citrate-stabilized spherical gold nanoparticles (diameter 11 nm, as determined by Differential Centrifugal Sedimentation, using a DC24000 disk centrifuge from CPS Ltd.) were synthesized following a modified Turkevich-Frens method.^60,61^ In brief, 20 mL of a hot 40 mM trisodium citrate solution in Milli-Q water were added to a boiling solution of 200 μmol HAuC1_4_ trihydrate in 200 mL Milli-Q water and refluxed under vigorous stirring for 60 min, the solution was then cooled overnight under stirring and filtered.

### CALNN-Capped Gold Nanoparticles

For GNP functionalization, CALNN peptide (Peptide Protein Research Ltd., used as received) was dissolved in Milli-Q water to give two stock solutions of 1 mg/mL and 0.05 mg/mL, respectively. To 0.5 mL of the GNP solution, 0.5 mL of Milli-Q water and 111.1 μL of the peptide stock solution were added and left overnight, the excess peptide was then removed by repetitive cycles of centrifugation and redispersion in fresh Milli-Q water, reducing the final volume to 0.5 mL.

### Sample Preparation for FTIR

In total, 18 mL of each GNP colloidal dispersion (using stock solutions with high and low CALNN concentration, respectively) were prepared. These were concentrated to a volume of 1 mL by lyophilisation and re-dispersion in Milli-Q water and cleaned by dialysis against Milli-Q water, using a 10 kDa cut-off membrane. For FTIR spectroscopy, the solvent was exchanged to D_2_C to avoid the strong H_2_O absorbance which overlaps with the amide I band; this was achieved by repeated cycles of lyophilisation and re-suspension in D_2_O. The GNP concentrations in the final samples were ~3 μM. as determined by UV-vis spectroscopy, using a sample diluted by a factor 400. A 10 mg/mL solution of CALNN in D_2_O was used to record the FTIR spectrum of free peptide.

### Fourier Transform Infrared Spectroscopy

FTIR spectra were recorded on a Bio-Rad FTS-40 FTIR spectrometer, averaging 1000 scans with 1 cm^−1^ resolution, using an IR cell with CaF_2_ windows and 50 μm spacer.

## ASSOCIATED CONTENT

### Supporting Information

Additional information on computational models with mobile or bound sulfur atoms; determination of CALNN capping density by FTIR; comments on the DSSP algorithm; construction and quantitative analysis of Ramachandran plots; polar charts of β-structure probability for CALNN and CFGAILSS monolayers; secondary structure analysis of CFGAILSS-capped 10 nm GNP simulated with different starting conditions; comparison of β-structure probability in CFGAILSS-capped 10 nm GNP simulated with the two computational models; distribution of backbone-backbone distances in CALNN-capped 25 nm GNP; temporal change of spatial β-structure distribution; movie of CFGAILSS-capped 10 nm GNP showing secondary structure assignment every ps over the last 100 ps of the 10 ns “production run” in AVI format. This material is available free of charge via the Internet at http://pubs.acs.org/.

Data access statement: The computational data and the custom-written software from this paper are available in an on-line repository of the University of Liverpool at DOI: 10.17638/datacat.liverpool.ac.uk/184

## ACKNOWLEDGMENTS

The authors acknowledge A*STAR Computational Resource Centre, the University of Liverpool and ARCHER UK National Supercomputing Service (http://www.archer.ac.uk) for the availability of high-performance computing resources. A.M.D. acknowledges support from EPSRC (DTP studentship). D.P. acknowledges support from A*STAR Joint Council Office (Grant number 14302FG094).

## Insert Table of Contents artwork here

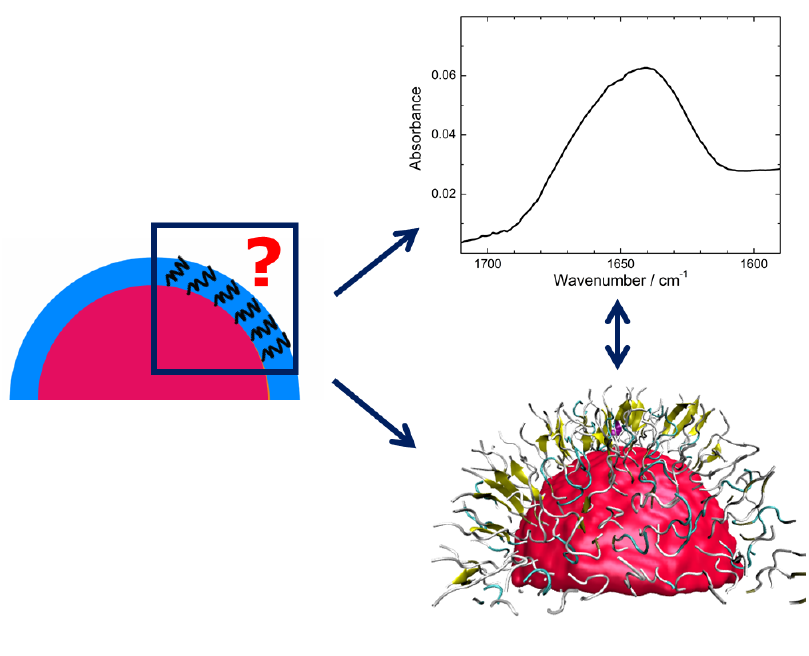

